# Whole genome sequencing-based detection of antimicrobial resistance and virulence in non-typhoidal *Salmonella enterica* isolated from wildlife

**DOI:** 10.1101/155192

**Authors:** Milton Thomas, Gavin John Fenske, Sudeep Ghimire, Ronald Welsh, Akhilesh Ramachandran, Joy Scaria

## Abstract

The aim of this study was to generate a reference set of *Salmonella enterica* genomes isolated from wildlife from the United States and to determine the antimicrobial resistance and virulence gene profile of the isolates from the genome sequence data. We sequenced the whole genomes of 103 *Salmonella* isolates sampled between 1988 and 2003 from wildlife and exotic pet cases that were submitted to the Oklahoma Animal Disease Diagnostic Laboratory, Stillwater, Oklahoma. Among 103 isolates, 50.48% were from wild birds, 0.9% was from fish, 24.27% each were from reptiles and mammals. 50.48% isolates showed resistance to at least one antibiotic. Resistance against the aminoglycoside streptomycin was most common while 9 isolates were found to be multi-drug resistant having resistance against more than three antibiotics. Determination of virulence gene profile revealed that the genes belonging to csg operons, the fim genes that encode for type 1 fimbriae and the genes belonging to type III secretion system were predominant among the isolates. The universal presence of fimbrial genes and the genes encoded by pathogenicity islands 1-2 among the isolates we report here indicates that these isolates could potentially cause disease in humans. Therefore, the genomes we report here could be a valuable reference point for future traceback investigations when wildlife is considered to be the potential source of human Salmonellosis.

## Background

*Salmonella enterica* is the leading cause of foodborne illness in the United States accounting for approximately 1.2 million infections, 23,000 hospitalizations and 450 deaths annually. Over the past few decades, *Salmonella* has acquired new virulence determinants that influence host-tropism which helps these organisms to adapt to a wide range of hosts1]. Multiple serovars of *S. enterica* originating from mammalian, reptilian and avian hosts have been reported to cause infections in humans[1]. Wildlife and exotic pets harboring *Salmonella* are potential sources for human infections[1]. Transmission of *Salmonella* from wildlife and exotic animals to humans occurs through multiple pathways. Increasing evidence suggests that there could be a bidirectional transmission of *Salmonella* between domesticated and wild animals. Farm animals acquiring *Salmonella* from wildlife, could increase the risk of human infection. *Salmonella* infections in humans have also been reported through direct contact with exotic pets and wildlife, especially those in captivity. Consumption of contaminated game bird meat is also a potential source for foodborne salmonellosis. Furthermore, wildlife such as rodents and birds, harboring in the proximity of food production units can act as carriers and contaminate food products leading to indirect infections.

The threat posed by salmonellosis is further compounded by the presence of resistance genes that confer resistance to multiple antimicrobial drugs. According to the National Antimicrobial Resistance Monitoring System (NARMS) integrated report, 20% of human *Salmonella* isolates exhibit antimicrobial resistance (AMR). Antimicrobial-resistant *Salmonella* infections result in increased disease severity and longer hospitalizations in addition to economic losses [2]. Research indicate that *Salmonella* isolates from various wildlife species also possess AMR determinants and the prevalence rate of AMR genes in these isolates could be as high as 100% [3, 4]. Thus, *Salmonella* in wildlife poses a significant risk to human health underlining the need for an integrative ‘One Health’ approach for the surveillance of pathogens among humans, domestic animals, and wildlife population.

Whole genome sequencing (WGS) of foodborne pathogens could be adopted as an effective and rapid surveillance tool. Compared to conventional antimicrobial tests, WGS offers a more comprehensive information on the genotypic characteristics of pathogens including identification of AMR and virulence determinants, and serotypes. Recent studies have utilized WGS to reliably predict the antimicrobial characteristics in various pathogens including *Salmonella* [5-8]. In this study, WGS was utilized to predict AMR and virulence determinants in *Salmonella* isolates from exotic pets and wildlife.

## Methods

### Quality assurance

All strains were identified as *Salmonella enterica* following the American Association of Veterinary Laboratory Diagnosticians certified laboratory. For genome sequencing, each isolate was streaked on Salmonella selective medium and a single colony was picked and used for further steps as outlined below.

### Salmonella bacterial isolates

A total of 103 *Salmonella* isolates were revived from archival cultures obtained from exotic pet or wildlife clinical specimens submitted to the Oklahoma Animal Disease Diagnostic Laboratory, Stillwater, Oklahoma during 1988 – 2003. The metadata for the samples used in this study are provided in Table 1. Isolates were streaked on Luria-Bertani agar slants and were transported to the Animal Disease Research and Diagnostic Laboratory, South Dakota State University, Brookings, South Dakota for WGS. Samples were streaked on Luria-Bertani agar plates upon arrival to the laboratory. A single bacterial colony from the agar plate was then inoculated to Luria-Bertani broth and cultured at 37^0^C.

**Table 1.** List of *Salmonella enterica* strains isolated and sequenced from wild life and the corresponding metadata.

| Strain ID | Serovar | Year | Animal | NCBI SRA BioSample ID | NCBI SRA ID |
| --- | --- | --- | --- | --- | --- |
| ADRDL-001 | poona | 1993 | Alligator Omentum | SAMN06330630 | SRR5278825 |
| ADRDL-002 | typhimurium | 1993 | Auodad Feces | SAMN06330629 | SRR5278822 |
| ADRDL-003 | gaminara | 1994 | Ratite Intestine | SAMN06330628 | SRR5278823 |
| ADRDL-004 | lille | 1993 | Gamebird Embryo | SAMN06330627 | SRR5278827 |
| ADRDL-005 | typhimurium | 1993 | Ratite Feces | SAMN06333495 | SRR5278819 |
| ADRDL-006 | typhimurium | 1993 | Ratite Feces | SAMN06333494 | SRR5278824 |
| ADRDL-007 | thompson | 1993 | Ratite Cecum | SAMN06333493 | SRR5278802 |
| ADRDL-008 | livington | 1993 | Ratite Cecum | SAMN06333492 | SRR5278806 |
| ADRDL-009 | typhimurium | 1993 | Ratite Feces | SAMN06333491 | SRR5278801 |
| ADRDL-010 | montevideo | 1993 | Ratite Feces | SAMN06333489 | SRR5278805 |
| ADRDL-011 | 6,7-nonmotile | 1993 | Ratite Intestine | SAMN06333488 | SRR5278804 |
| ADRDL-012 | arechavaleta | 1994 | Ratite Intestine | SAMN06333486 | SRR5278803 |
| ADRDL-013 | 4,5,12:i-monophasic | 1994 | Ratite Liver | SAMN06333485 | SRR5380966 |
| ADRDL-014 | berta | 1994 | Ratite Intestine | SAMN06333484 | SRR5278808 |
| ADRDL-015 | ituri | 1994 | Ratite Cecum | SAMN06333483 | SRR5278773 |
| ADRDL-016 | ituri | 1994 | Ratite Intestine | SAMN06333482 | SRR5278772 |
| ADRDL-017 | heidelberg | 1993 | Wild Turkey Liver | SAMN06333481 | SRR5278779 |
| ADRDL-018 | heidelberg | 1993 | Wild Turkey Liver | SAMN06333480 | SRR5278777 |
| ADRDL-019 | godesberg | 1993 | Wild Turkey Cecum | SAMN06333479 | SRR5278778 |
| ADRDL-020 | 4,5,12:i-monophasic | 1993 | Eclectus Colon | SAMN06333477 | SRR5278771 |
| ADRDL-021 | anatum | 1993 | Giraffe Feces | SAMN06333476 | SRR5278774 |
| ADRDL-022 | anatum | 1993 | Giraffe Feces | SAMN06333475 | SRR5278780 |
| ADRDL-023 | pomona | 1993 | Python Abdominal swab | SAMN06333473 | SRR5278767 |
| ADRDL-024 | muenchen | 1993 | Ratite Intestine | SAMN06333472 | SRR5278776 |
| ADRDL-025 | typhimurium | 1994 | Rodent Intestine | SAMN06333471 | SRR5278770 |
| ADRDL-026 | hadar | 1995 | Wild Chicken Intestine | SAMN06333470 | SRR5278768 |
| ADRDL-027 | hadar | 1994 | Ratite Intestine | SAMN06333469 | SRR5278769 |
| ADRDL-028 | typhimurium | 1988 | Primate Intestine | SAMN06333465 | SRR5278873 |
| ADRDL-029 | albany | 1988 | Saiga Intestine | SAMN06333464 | SRR5278882 |
| ADRDL-030 | arizona | 1988 | Snake | SAMN06333462 | SRR5330438 |
| ADRDL-031 | arizona | 1989 | Boa Intestinal swab | SAMN06333460 | SRR5330446 |
| ADRDL-032 | 16:z10-e,n,xz15 | 1989 | Cervine Feces | SAMN06333459 | SRR5330441 |
| ADRDL-033 | enteritidis | 1989 | Hedgehog Spleen | SAMN06333458 | SRR5330440 |
| ADRDL-034 | typhimurium(O5-)* | 1992 | Pigeon Airsac swab | SAMN06333457 | SRR5330448 |
| ADRDL-035 | typhimurium | 1989 | Screech owl Liver | SAMN06333455 | SRR5330445 |
| ADRDL-036 | braenderup | 1989 | Snow leopard Intestine | SAMN06333454 | SRR5330444 |
| ADRDL-037 | saintpaul | 1989 | Snow leopard Lung | SAMN06333453 | SRR5330406 |
| ADRDL-038 | montevideo | 1992 | Cervid Intestine | SAMN06333451 | SRR5329403 |
| ADRDL-039 | enteritidis | 1993 | Emu Feces | SAMN06333450 | SRR5329404 |
| ADRDL-040 | enteritidis | 1993 | Emu Feces | SAMN06333449 | SRR5380965 |
| ADRDL-041 | worthington | 1992 | Quail Intestine | SAMN06333448 | SRR5380958 |
| ADRDL-042 | II 43:z4,z23:- or IIIa 43:z4,z23:- or Farmingdale or IV 43:z4,z23:-* | 1992 | Reptile Eggsac | SAMN06333447 | SRR5329405 |
| ADRDL-043 | panama | 1992 | Rhea Intestine | SAMN06333694 | SRR5409894 |
| ADRDL-044 | ituri | 1994 | Ratite Cecum | SAMN06333692 | SRR5409893 |
| ADRDL-045 | newport | 1995 | Ratite Feces | SAMN06333691 | SRR5409493 |
| ADRDL-046 | newport | 1995 | Dolphin Lung | SAMN06333689 | SRR5409890 |
| ADRDL-047 | typhimurium | 1997 | Psittacine Lung | SAMN06333684 | SRR5409485 |
| ADRDL-048 | typhimurium | 1997 | Psittacine Intestine | SAMN06333683 | SRR5409315 |
| ADRDL-049 | muenchen | 1996 | Ratite Intestine | SAMN06333682 | SRR5409313 |
| ADRDL-050 | schwazengrund | 1997 | Ratite Intestine | SAMN06333681 | SRR5409312 |
| ADRDL-051 | archavaleta | 1997 | Antelope Intestine | SAMN06333692 | SRR5409893 |
| ADRDL-052 | infantis | 1997 | Fish Water | SAMN06645614 | SRR5398012 |
| ADRDL-053 | bredeney | 1998 | Llama Intestine | SAMN06333861 | SRR5409360 |
| ADRDL-054 | plymouth | 1997 | Reptile Liver | SAMN06330627 | SRR5278827 |
| ADRDL-055 | montevideo | 1997 | Reptile Intestine | SAMN06645663 | SRR5398013 |
| ADRDL-056 | branderup | 1995 | Wild Chicken Intestine | SAMN06645590 | SRR5387496 |
| ADRDL-057 | enteriditis | 1996 | Wild Chicken Intestine | SAMN06645569 | SRR5387492 |
| ADRDL-058 | typhimurium | 1996 | Wild Chicken Feces | SAMN06645567 | SRR5387491 |
| ADRDL-059 | bredeney | 1995 | Gamebird Intestine | SAMN06645592 | SRR5387497 |
| ADRDL-060 | livingston | 1996 | Gamebird Intestine | SAMN06645590 | SRR5387496 |
| ADRDL-061 | enteriditis | 1995 | Psittacine Intestine | SAMN06645588 | SRR5387490 |
| ADRDL-062 | montevideo | 1996 | Psittacine Liver | SAMN06645587 | SRR5387493 |
| ADRDL-063 | 7,14:K-monophasic | 1995 | Ratite Intestine | SAMN06645585 | SRR5387523 |
| ADRDL-064 | anatum | 1995 | Ratite Feces | SAMN06645654 | SRR5387521 |
| ADRDL-065 | enteriditis | 1995 | Ratite | SAMN06645582 | SRR5387527 |
| ADRDL-066 | thompson | 1995 | Ratite Cloacal swab | SAMN06645594 | SRR5387519 |
| ADRDL-067 | thompson | 1995 | Ratite Cloacal swab | SAMN06645593 | SRR5387517 |
| ADRDL-068 | 4,5,12: i | 1995 | Ratite Pericardial fluid | SAMN06645652 | SRR5387518 |
| ADRDL-069 | livingston | 1996 | Llama Intestine | SAMN06645650 | SRR5387514 |
| ADRDL-070 | uganda | 1999 | Cervine Intestine | SAMN06645664 | SRR5398014 |
| ADRDL-071 | lille | 2000 | Cervine Intestine | SAMN06645663 | SRR5398013 |
| ADRDL-072 | parera | 1998 | Iguana Cloacal swab | SAMN06645662 | SRR5398016 |
| ADRDL-073 | anatum | 1998 | Ratite Feces | SAMN06645661 | SRR5398025 |
| ADRDL-074 | anatum | 1998 | Ratite Feces | SAMN06645615 | SRR5398018 |
| ADRDL-075 | kiambu | 1998 | Ratite Cloacal swab | SAMN06645614 | SRR5398012 |
| ADRDL-076 | marina | 2000 | Reptile Feces | SAMN06645660 | SRR5398017 |
| ADRDL-077 | bredeney | 2003 | Alpaca Liver | SAMN06645613 | SRR5398015 |
| ADRDL-078 | sandiego | 2003 | Alpaca Feces | SAMN06645612 | SRR5398009 |
| ADRDL-079 | sandiego | 2003 | Alpaca Feces | SAMN06645611 | SRR5398010 |
| ADRDL-080 | bredeney | 2003 | Antelope Feces | SAMN06645610 | SRR5398011 |
| ADRDL-081 | virginia or muenchen* | 2002 | Ratite | SAMN06645609 | SRR5398008 |
| ADRDL-082 | newport* | 2002 | Ratite | SAMN06645659 | SRR5398007 |
| ADRDL-083 | enteritidis* | 2002 | Ratite | SAMN06645658 | SRR5398001 |
| ADRDL-084 | oranienburg | 2003 | Iguana Cloacal swab | SAMN06645657 | SRR5398006 |
| ADRDL-085 | give | 2003 | Iguana Cloacal swab | SAMN06658957 | SRR5409330 |
| ADRDL-086 | chameleon | 2003 | Iguana Cloacal swab | SAMN06333875 | SRR5387539 |
| ADRDL-087 | typhimurium | 2002 | Llama Feces | SAMN06333874 | SRR5387538 |
| ADRDL-088 | anatum | 2003 | Llama Feces | SAMN06333873 | SRR5387533 |
| ADRDL-089 | typhimurium | 2003 | Llama Feces | SAMN06333872 | SRR5387534 |
| ADRDL-090 | agona | 2003 | Marsupial Intestine | SAMN06333871 | SRR5387532 |
| ADRDL-091 | miami | 2001 | Reptile Fecal swab | SAMN06658960 | SRR5409328 |
| ADRDL-092 | arizona | 2001 | Reptile Liver | SAMN06658959 | SRR5409327 |
| ADRDL-093 |  | 2001 | Reptile Cloacal swab | SAMN06658958 | SRR5409325 |
| ADRDL-094 | marina | 2002 | Reptile Cloacal swab | SAMN06658962 | SRR5409322 |
| ADRDL-095 | marina | 2002 | Reptile Abscess swab | SAMN06658961 | SRR5409324 |
| ADRDL-096 | arizona | 2002 | Reptile Lung | SAMN06333869 | SRR5387526 |
| ADRDL-097 | parera | 2002 | Reptile Cloacal swab | SAMN06333866 | SRR5397979 |
| ADRDL-098 | chameleon | 2002 | Reptile Cloacal swab | SAMN06333865 | SRR5397978 |
| ADRDL-099 | senftenberg | 2002 | Reptile Cloacal swab | SAMN06333864 | SRR5397977 |
| ADRDL-100 | arizona | 2002 | Reptile Cloacal swab | SAMN06333863 | SRR5409363 |
| ADRDL-101 | arizona | 2002 | Reptile Cloacal swab | SAMN06333862 | SRR5409361 |
| ADRDL-102 | kisarwe | 2003 | Reptile Cloacal swab | SAMN06333861 | SRR5409360 |
| ADRDL-103 | newport | 2003 | Turtle Intestine | SAMN06333859 | SRR5409359 |

\* Predicted serovar using SeqSero

### Genomic DNA isolation and WGS

Genomic DNA was isolated from 1.0 mL overnight cultures using the Qiagen DNeasy kits (Qiagen, Valencia, CA, USA) according to manufacturer’s protocol. The quality of isolated DNA was analyzed using NanoDrop™ One (Thermo Scientific™, DE) and was quantified using Qubit^®^ 3.0 (Thermo Fisher Scientific Inc., MA) fluorometer and stored at -20^0^C until use. Whole-genome sequencing was performed on Illumina Miseq platform using V2 chemistry with 2 X 250 paired-end chemistry Briefly, the concentrations of genomic DNA samples were adjusted to 0.3 ng/μl concentration and were processed using Nextera XT DNA Sample Prep Kit (Illumina Inc. San Diego, CA). The libraries were normalized using bead-based procedure and pooled together at equal volume. The pooled library was denatured and sequenced using Miseq reagent version 2 (Illumina, Inc., CA).

### Genome Assembly and Identification of Resistance and Virulence Genes

The raw data files were de-multiplexed and converted to FASTQ files using Casava v.1.8.2. (Illumina, Inc, San Diego, CA). The FASTQ files were trimmed and assembled *de novo* using CLC workbench 9.4 (Qiagen Bioinformatics, CA). The antibiotic resistance genes in the assembled *Salmonella* genomes were identified by BLAST search against a local copy of the antibiotic resistance gene sequence data from ResFinder [9] and CARD[10]. The parameters used for BLAST search were ≥ 95% gene identity and 50% sequence length of the resistance gene. The virulence genes in the genomes were predicted using a similar approach. *Salmonella* virulence gene sequences were extracted from Virulence Factor Database[11] and *Salmonella* genome assemblies were searched against these sequences using BLAST with ≥ 90% gene identity and 50% sequence length cut off.

### Serotyping and antimicrobial susceptibility test

Serotypes of the strains were determined at the National Veterinary Service Laboratory, Ames, IA. Antimicrobial susceptibility of all *Salmonella* isolates was determined using the Sensititre NARMS Gram Negative Plate (CMV3AGNF, Thermofisher). The antibiotics used were gentamicin, streptomycin, amoxicillin-clavulanic acid, ampicillin, cefoxitin, ceftiofur, ceftriaxone, azithromycin, chloramphenicol, nalidixic acid, ciprofloxacin, sulfisoxazole, trimethoprim-sulfamethoxazole, and tetracycline. The AMR was determined according to Clinical and Laboratory Standards Institute guidelines except for azithromycin and sulfisoxazole where the data obtained was indeterminate and were not included in further analysis.

## Results and Discussion

### Distribution of Salmonella isolates among wildlife and exotic pets

A total of 103 *Salmonella* isolates sampled between 1988 and 2003 from wildlife and exotic pets were included in the present study for determining the antimicrobial susceptibility using whole genome sequencing. Among 103 isolates, 52 isolates (50.48%) were from wild birds, 1 isolate (0.9%) was from fish, 25 isolates each (24.27%) were from reptiles and mammals (Table 1). The serovars of 96 isolates in this study were determined at the National Veterinary Service Laboratory, Ames, IA, and the remaining 6 serovars were predicted using Seqsero [12]. The serovar of one isolate (ADRDL-093) was not identified under Kauffmann-White classification. A total of 45 serovars were identified among the 103 isolates, of which Typhimurium (12.62%) was the most frequent serovar. Other serovars that had higher prevalence were Enteritidis (6.8%), Anatum (5.8%), Arizona (5.8%), Bredeney (3.9%) and Montevideo (3.9%). The presence of multiple serotypes in wildlife has also been reported from previous epidemiological studies. 9 *Salmonella* samples isolated from marine mammals and birds in California yielded 7 serovars [4]. Similar to our findings, *Salmonella* Typhimurium was reportedly the predominant serovar present in wildlife [13-15] in various parts of the world.

### Phenotypic resistance to antimicrobials

Antimicrobial susceptibility test of 103 *Salmonella* bacterial isolates was performed using Sensititre NARMS gram-negative plate. The results were classified into 3 categories: resistant, intermediate or susceptible. 52 out of the 103 isolates (50.48%) showed resistance to at least one antibiotic (Fig 1a). Resistance against the aminoglycoside streptomycin was most commonly observed. 48 of the 103 isolates (46.6%) exhibited this phenotype. However, only three isolates (2.9%) showed resistance against gentamicin which also belongs to the aminoglycoside class of antibiotics. The isolates with resistance against gentamycin were also resistant to streptomycin. In the beta-lactam group, ampicillin resistance was the most common phenotype and was seen in 11 of the isolates (10.67%). Among these 11 isolates, few also shared resistance against other beta-lactams such as amoxicillin-clavulanic acid (4), cefoxitin (3), and ceftiofur (3). All the isolates were susceptible to ceftriaxone except one with intermediate resistance. The isolates that were susceptible to ampicillin were also susceptible to all other beta-lactams.

**Figure 1.**
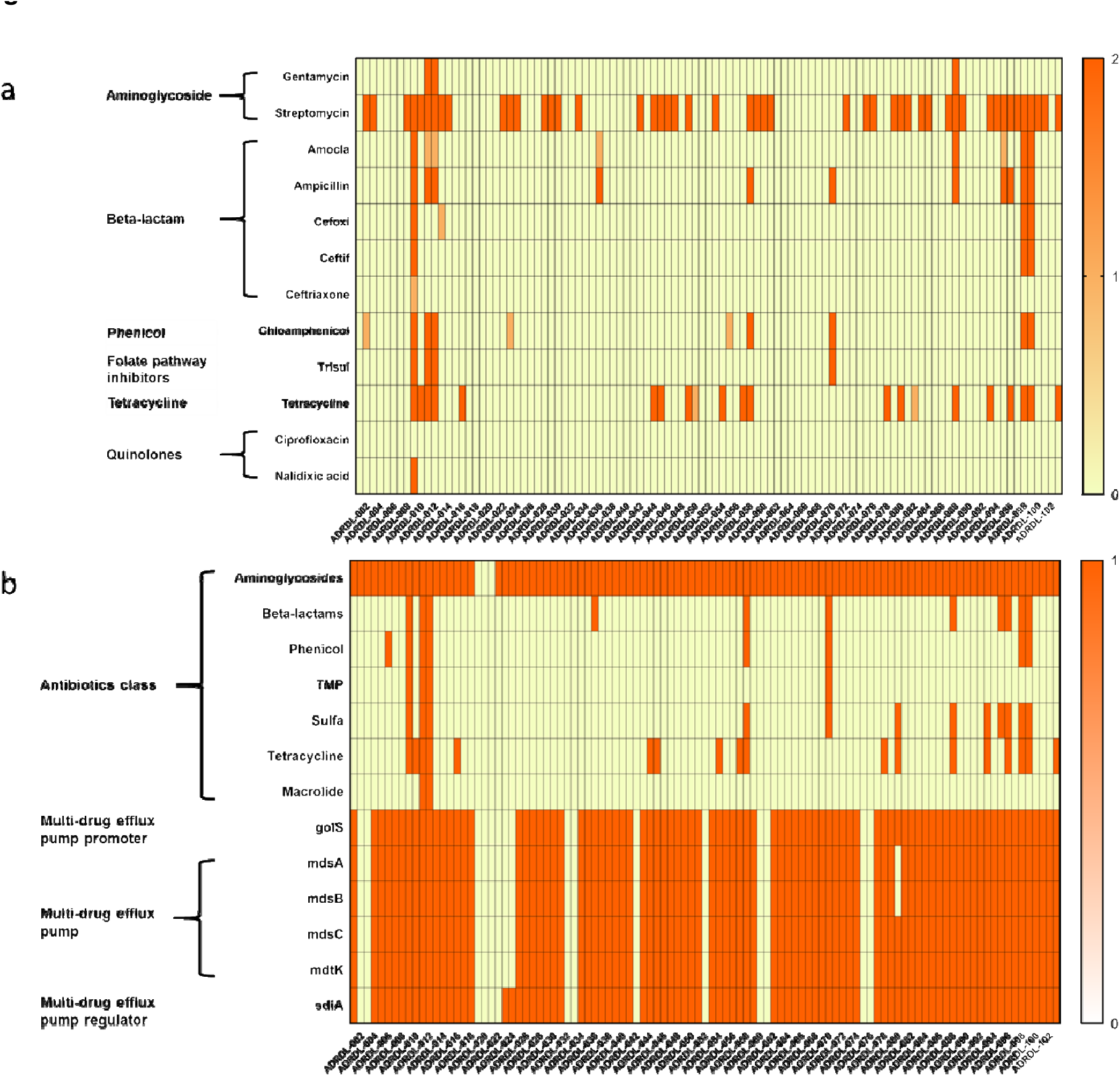
Phenotypic and genotypic anti-microbial resistance of 103 wildlife salmonella isolates. Fig1a - Heatmap of phenotypic resistance against 12 antibiotics was measured using Sensititre NARMS gram-negative plate. The CLSI breakoff points for resistance against various antibiotics was used for determining the antimicrobial susceptibility of 103 salmonella strains. Legend description: 0 = susceptible, 1 = intermediate, and 2 = resistant. Complete data used to generate Fig1 B is given in additional tables 1. Fig1b - Heatmap of genotypic resistance against antimicrobials detected using CLC workbench 9.0 by BLAST against ResFinder 2.1 and CARD database. Legend description: 0 = absent, 1 = present. The golS, mdsABC complex, and mdtK genes associated with multidrug resistance was present in all except 16 isolates. Similarly, multidrug efflux pump regulator gene sdiA was absent in 14 isolates. Data underlying this figure is given in Supplemental file 1.

Chloramphenicol resistance was observed for seven isolates (6.7%), trimethoprim-sulfamethoxazole resistance in 4 (3.88%) and tetracycline resistance in 19 (18.44%) of the isolates. All the isolates were susceptible to ciprofloxacin and all except one isolate was susceptible to nalidixic acid. Nine isolates were found to be multi-drug resistant having resistance against more than three antibiotics.

### Genotypic resistance to antimicrobials

The presence of genes that could contribute to AMR was detected by BLAST searching the assembled *Salmonella* genomes against a local copy of Resfinder and CARD sequence data (Fig 1b). Bacterial isolates showing “intermediate” resistance on antimicrobial susceptibility test was grouped with “susceptible” isolates for the calculation of sensitivity and specificity of AMR genotype. 22 genes were detected which provide resistance to aminoglycosides and the genes were present in 100 isolates. The sensitivity was 100% and specificity was 5.45 % for resistance against aminoglycosides. The low specificity was probably due to the lack of resistance genes being expressed *in vitro*. Genes responsible for resistance to betalactam antibiotics were detected in 11 isolates which were also resistant by antimicrobial susceptibility test. The plasmid-mediated cephalosporinase gene blaLAT-1, plasmid-borne class C beta-lactamase gene blaBIL-1, and blaCMY (Class C) genes were found together and were detected in three isolates. Genes belonging to blaTEM (class A) were found in eight isolates. Collectively, there were 280 beta-lactamase genes present in those 11 isolates. The sensitivity and specificity was 100 % for beta-lactams. Phenicol resistance encoded by cat, catA1 and floR genes was present in 8 isolates. The sensitivity was 100% and specificity was 98.96 % for phenicol resistance. dfrA1, dfrA10, dfrA12, sul1, sul2, and sul3 genes conferring resistance to trimethoprim-sulfamethoxazole drugs were present in 12 isolates. The sensitivity was 100% and specificity was 91.92 % for trimethoprim-sulfamethoxazole. The sul1, sul2, and sul3 genes could also contribute to resistance against sulfisoxazole. However, a definite conclusion of genotype-phenotype correlation is lacking due to the absence of antimicrobial susceptibility test data that matches the CLSI recommended break point for resistance against sulfisoxazole. Tetracycline resistance encoded by tet(A), tet(B), tet(C), and tet(D) genes for tetracycline efflux pumps were detected in 18 samples all of which were also resistant by antimicrobial susceptibility test. The sensitivity was 94.74% and specificity was 100% for tetracycline resistance. Two isolates carried the mph(A) gene which confers resistance to macrolides. However, the only macrolide that was tested was azithromycin and the genotype-phenotype relation could not be established due to lack of data from antimicrobial susceptibility test that matches with the break point recommended by CLSI (> 32 mg/L).

Overall, the sensitivity for detecting AMR using genotype was 100% except for tetracycline where 1 isolate was phenotypically resistant even in the absence of the (tet) gene. The specificity for aminoglycosides had the highest degree of incongruence between genotype and phenotype. 52 isolates that were positive for aminoglycoside resistance genes were phenotypically susceptible. Although not to the degree found in this study, a mismatch in phenotype-genotype correlation was also reported previously in *E. coli* and *Salmonella* for aminoglycoside resistance, especially for streptomycin [5, 16]. There was 100% phenotype-genotype correlation for beta-lactam resistance. Phenicols and tetracycline also had >98% specificity, while trimethoprim-sulfamethoxazole had lower specificity (91.2%) because of four isolates that were genotypically resistant but phenotypically susceptible. These results are also similar to those obtained in previous studies [5, 16] where correlation approaching 100% was obtained for antimicrobials other than aminoglycosides.

In addition to the genes that confer AMR, we also analyzed the genes that could confer multi-drug resistance (Fig 1b). The golS gene is a promoter for multidrug efflux pump, mdsABC [17] and was detected among 84.46% (n=87) isolates. Similarly, mdsABC (multidrug transporter of *Salmonella*) complex which is made up of mdsA, mdsB, and mdsC units, was found in all isolates that had golS gene except one isolate which lacked mdsB and mdsC genes. The mdsABC complex is known to provide resistance against a variety of drugs and toxins and is involved in Salmonella virulence and pathogenicity [17, 18]. The mdtK gene, a multi-efflux pump which could provide resistance against norfloxacin, doxorubicin and acriflavin [18] and sdiA, a regulator for multi-drug resistance pump AcraB [19], were present in 84.46% and 86.41% of the isolates respectively. The presence of these genes could contribute to the virulence and pathogenicity of these *Salmonella* isolates and also indicates the potential for these isolates to resist various antibiotics and toxins.

### Analysis of virulence determinants

The genes that are associated with virulence among 103 wildlife *Salmonella* isolates were analyzed (Fig 2) using CLC workbench 9.4. The parameters used were the minimum identity of 90% and minimum length of 50%. A total of 197 virulence genes were detected by BLAST search against a local copy of the Virulence Factor Database. The virulence-associated determinants collectively were grouped under 9 categories: fimbrial adherence determinants, macrophage inducible genes, determinants associated with magnesium uptake, nonfimbrial adherence determinants, genes associated with secretion system, serum resistance determinants, stress proteins, toxins, and two-component regulatory systems.

**Figure 2.**
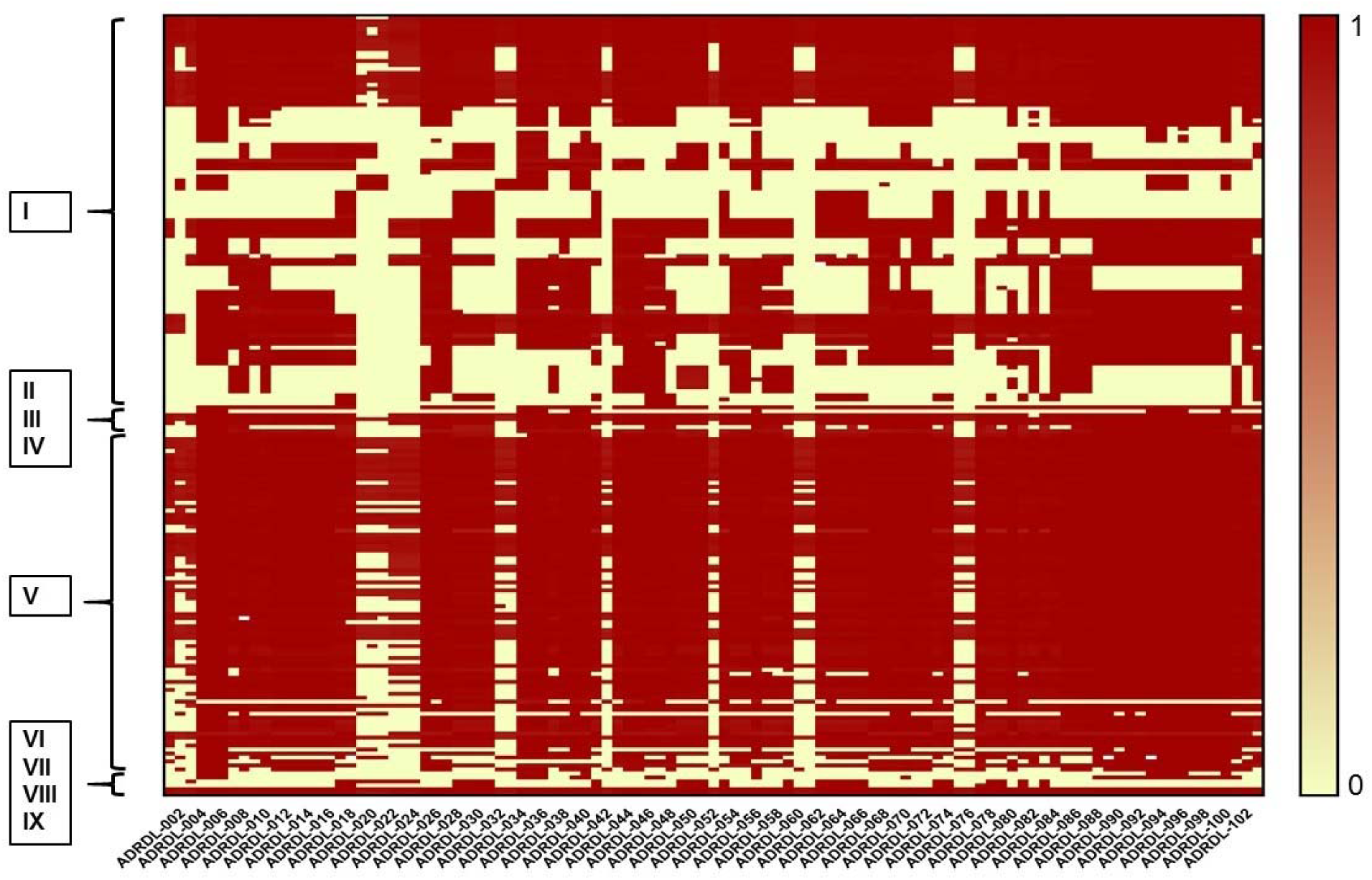
Heatmap of virulence genes present in 103 wildlife salmonella isolates. Salmonella genome Assemblies were searched against a local copy of the Virulence Factor Database using BLAST. In the figure, each row represents a virulence gene and each column denotes a sample. Legend on the left side of the figure denotes the following categories of virulence genes: (I) Fimbrial adherence determinants, (II) Macrophage inducible genes, (III) Magnesium uptake, (IV) Non-fimbrial adherence determinants, (V)Secretion system, (VI) Serum resistance, (VII)Stress protein, (VIII)Toxin, and (IX) Two-component system. The virulence genes belonged 5 categories – PSLT, SeHA, SEN, SeSA, and STM. Seventeen isolates had fewer virulence genes compared to others and this correlated with the absence of genes associated with multidrug resistance. Legend description: 0 = absent, 1 = present. Data underlying this figure is given in Additional table 2 .

Among fimbrial adherence determinants, the genes belonging to two csg operons csgBAC and csgDEFG were present universally in all isolates. These genes encode for curli fimbriae or thin aggregative fimbriae and mediate binding to various serum and tissues matrix proteins [20]. Another gene cluster that was ubiquitously present were the *fim* genes that encodes for type 1 fimbriae. This cluster is comprised of the *fimAICDHF* operon and three regulatory genes *fimW, fimY,* and *fimZ* and mediates adherence to eukaryotic cells [21]. However, the *fimY* gene was not detected in ten isolates at the BLAST search cut-off level we used.

The genes belonging to type III secretion system (TTSS/T3SS) encoded by Salmonella pathogenicity island -1 (SPI-1) and -2 (SPI-2) were also predominantly present among the isolates. This included SPI-1 regulator genes *hilACD*, and SPI-1 encoded *inv/spa*, and *prg/org* operons that were detected in all the isolates. Similarly, SPI-2 regulatory gene ssrB, chaperone protein encoding genes - sscA and sscB, and ssa genes that encode for T3SS2 apparatus were also present among 103 isolates. However, the sse genes which encodes for the effectors were observed only in fewer isolates. Another set of genes that were present in all isolates were the genes that respond to magnesium level in the extracellular environment [22]. This included *mgtc,* which mediates magnesium uptake and *phoP-phoQ* genes that are regulators of the two-component system.

The least abundant virulence determinants were the *tcf, sta,* and *pef* fimbrial operons and *spv* gene cluster. These genes belonging to the fimbrial adherence determinants category were detected in less than 25% of the isolates. Additionally, *rck* gene that provides protection against the complement-mediated immune response of the host was also found in low abundance. There were 16 isolates that possessed fewer than 50% of the total virulence genes in the database (Fig 2). These isolates include ADRDL-002, -003, -019, -020, -021, -022, -023, -024, -032, -033, -042, -052, -060, -061, -075, and -076. Importantly, these isolates also had a lower abundance of genes that contributes to multi-drug resistance (Fig 1). However, these isolates come under various serotypes and were isolated from different host species. Therefore, a common factor responsible for the observed low abundance of virulence genes is not evident. The universal presence of fimbrial genes and the genes encoded by pathogenicity islands 1- 2 among the isolates we report here indicates that these isolates could potentially cause disease in humans. Therefore, the genomes we report here could be a valuable reference point for future traceback investigations in instances where wildlife may be considered as a potential source of human Salmonellosis.

## List of abbreviations

AMR: Antimicrobial Resistance
NARMS: The National Antimicrobial Resistance Monitoring System
WGS: Whole Genome Sequencing

## Declarations

### Acknowledgements

Authors thank the Section of Bacteriology, Animal Disease Research and Diagnostic Laboratory, South Dakota for helping with the antimicrobial susceptibility testing of the *Salmonella* isolates. We also thank Scott Talent and Leanne Tillman (Oklahoma Animal Disease Diagnostic Laboratory, Stillwater, OK) for reviving archival cultures necessary for this study.

### Author Contributions

JS and AR conceived and designed the study. MT, GJF and SG performed the experiments. RW originally developed the culture archive. MT analyzed the data. MT, JS and AR wrote the manuscript. All authors read and approved the final manuscript.

### Availability of data and material

Genome sequence data of 103 *Salmonella enterica* isolates have been submitted in NCBI Sequence Read Archive (NCBI SRA) for public access. NCBI SRA accession number for 103 isolates described in this manuscript is given in Table 1.

---

**Additional Table 1: Genotypic Antibiotic resistance gene profile of 103 Salmonella isolates.** Salmonella genome assemblies were searched against a local copy of the Resfinder database using BLAST. Cut off parameter for BLAST search was ≥ 95% gene identity and 50% sequence length of the resistance gene. Values in the sample columns indicate the BLAST sequence percentage identity cutoff values.

**Additional Table 2: Mapping of virulence genes present in 103 wildlife salmonella isolates.** Salmonella genome assemblies were searched against a local copy of the Virulence Factor Database using BLAST. Minimum identity of 90% and minimum length of 50% BLAST hits were used as cut off value. Reference column indicates the NCBI gene locus tag of the reference genes used. Values in the sample columns indicate the BLAST sequence percentage identity cutoff values.

